# FM-GPT: Bayesian fine mapping for phenome-wide transcriptome-wide association studies

**DOI:** 10.64898/2026.04.07.717122

**Authors:** Travis Canida, Zhenyao Ye, Shao-Hsuan Wang, Hsin-Hsiung Huang, Yezhi Pan, Menglu Liang, Shuo Chen, Tianzhou Ma

## Abstract

Transcriptome-wide association studies (TWAS) integrate genome wide association studies with expression quantitative trait locus reference panels to identify genes associated with traits of interest. However, linkage disequilibrium and correlated gene expression can induce spurious TWAS signals, motivating fine mapping methods to prioritize putatively causal genes within associated loci. The rapid growth of large-scale phenomic resources (e.g. electronic health records (EHRs)) has shifted genetic studies from single-trait analyses to phenome-wide investigations that jointly evaluate many closely related phenotypes. We introduce FM-GPT (**F**ine-**m**apping of causal **G**enes for **P**henome-wide **T**ranscriptome-wide association studies), a novel Bayesian fine mapping method for prioritizing causal genes across multiple correlated phenotypes with potentially mixed outcome types (e.g., binary, count or continuous) in phenome-wide TWAS. FM-GPT performs gene-guided dimension reduction of the phenotypes and reveals pleiotropic or phenotype-specific effects of the identified genes. In simulations, FM-GPT identified true causal genes more accurately than other fine mapping methods while controlling false positives. We applied FM-GPT to two applications using data from UK Biobank: a brain-wide genetic analysis of MRI data derived regional cortical thickness measures and a phenome-wide genetic analysis of clinical phenotypes derived from EHR data. FM-GPT greatly narrowed down the set size of putatively causal genes and identified: 1. genes with pleiotropic effects on regional cortical thickness across the cerebral cortex, including five genes *BCAS3, LRRC37A, NOS2P3, ARL17B* and *UBB* on chromosome 17 regulating neuronal morphology and cortical organization; and 2. genes that influence multiple medical conditions across the circulatory, metabolic, digestive, respiratory and genitourinary systems, revealing two major axes of variation among these conditions that point to a potential trade-off in gene regulation between immune and metabolic functions. These results highlight FM-GPT’s power to disentangle complex gene–phenotype relationships in large-scale phenome-wide studies, uncovering shared biological mechanisms across diverse human traits and advancing translational and comorbidity research.

**Author Summary:** We developed a novel fine mapping method called FM-GPT, to identify putatively causal genes from correlated noise influencing a wide range of human traits and diseases with potentially mixed outcome types (e.g., binary, count or continuous). The rapid expansion of large-scale phenomic datasets has shifted the single-trait genetic studies to phenome-wide analyses, enabling the study of genetic architecture across many related traits simultaneously. FM-GPT performs gene-guided dimension reduction of the phenotypes and reveals pleiotropic or phenotype-specific effects of the identified causal genes. When applied to the UK Biobank data, FM-GPT greatly narrowed down the set size of putatively causal genes compared to other methods. The tool identified genes with pleiotropic effects on regional cortical thickness that regulate neuronal morphology and cortical organization across the cerebral cortex. It also identified genes that influence multiple medical conditions spanning the circulatory, metabolic, digestive, respiratory, and genitourinary systems. Among these conditions, two major axes of variation emerged, revealing a potential trade-off in gene regulation between immune and metabolic functions. This work provides a clearer picture of shared biological mechanisms across traits and diseases, advancing translational research and the understanding of comorbidity.

## Introduction

Transcriptome Wide Association Studies (TWAS) integrate Genome Wide Association Studies (GWAS) with expression quantitative trait locus reference panels (e.g. GTEx(1)) to identify genes whose genetically regulated expressions (GReX) are associated with a trait of interest(2, 3). TWAS typically proceed in two steps: First, gene expression prediction models are trained using reference panels with matched genotype and gene expression data, and these models are then used to impute GReX in GWAS. Second, association analyses are performed between GReX and the trait of interest to identify candidate susceptibility genes. Since the introduction of the first TWAS method PrediXcan(4), many TWAS methods have been developed that employ different statistical learning methods for expression prediction(5-7), integrate information across multiple tissues(8-10), or leverage GWAS summary statistics(8, 11, 12). Despite these advances, interpreting TWAS associations remains challenging: TWAS methods test genes individually without accounting for the correlation among predicted gene expressions arising from linkage disequilibrium (LD) among variants or shared regulatory architecture(2, 3, 13). To address this issue, several TWAS fine mapping methods (14-16) have been developed to prioritize putatively causal gene within associated regions. However, existing TWAS fine-mapping methods are primarily designed for single-trait analyses, limiting their applicability in settings where multiple related phenotypes are analyzed jointly.

Meanwhile, the rapid expansion of large-scale phenomic datasets has shifted the focus of human genetics from single-trait studies to phenome-wide analyses. Resources such as the UK Biobank (UKB) provide extensive collections of phenotypes derived from electronic health records (EHR), imaging-derived phenotypes (IDPs) and other clinical or non-clinical measurements. Phenome-wide TWAS analyses offer an opportunity to systematically investigate how genes regulate a broad spectrum of phenotypic outcomes. However, these analyses also introduce new methodological challenges. First, phenotypes in large-scale phenomic datasets are often highly correlated, and analyzing them independently can substantially increase the multiple-testing burden while reducing statistical power. Secondly, phenome-wide datasets frequently contain mixed outcome types, including continuous, categorical, and count outcomes, which complicates joint modeling(17-19). Thirdly, gene-phenotype relationships are inherently complex: individual genes may influence multiple phenotypes (pleiotropy), while most phenotypes are regulated by multiple genes (polygenicity). Disentangling these many-to-many relationships across both the transcriptome and the phenome remains an important and largely unresolved challenge.

To fill the gap, we propose FM-GPT (Fine-Mapping of causal Genes for Phenome-wide Transcriptome-wide association studies), a novel Bayesian fine mapping method for identifying causal genes across a number of related phenotypes in phenome-wide TWAS. FM-GPT jointly prioritizes causal genes for multiple correlated phenotypes while accommodating outcomes of mixed data types (e.g., continuous, binary or count). The method leverages GReX information to guide dimension reduction of the phenotypes, prioritizes putatively causal genes associated with the latent phenotype factors, and reveals pleiotropic or phenotype-specific effects of the identified genes. FM-GPT enables computationally scalable analysis of large phenomic datasets while facilitating the interpretation of complex gene-phenotype relationships and possibly heterogeneous pleiotropic effects.

Through extensive simulations, we demonstrate that FM-GPT improves the identification of true causal genes relative to existing TWAS fine-mapping approaches while controlling false positives. We further applied FM-GPT to two real data examples in the UK Biobank (UKB). In an brain-wide genetic analysis of MRI data derived regional cortical thickness measures, FM-GPT greatly narrowed down the set size of putatively causal genes compared to other fine mapping methods and identified a set of causal genes with pleiotropic effects on regional cortical thickness across the cerebral cortex, including five genes *BCAS3, LRRC37A, NOS2P3, ARL17B* and *UBB* on chromosome 17 regulating neuronal morphology and cortical organization. In a phenome-wide genetic analysis of EHR data derived clinical phenotypes, we identified genes that influence multiple medical conditions across the circulatory, metabolic, digestive, respiratory and genitourinary systems, and revealed two major axes of variation among these conditions pointing to a potential trade-off in gene regulation between immune and metabolic functions. These results highlight the utility of FM-GPT for disentangling complex gene-phenotype relationships in large-scale phenome-wide genetic studies, enabling the identification of shared biological mechanisms across related traits and diseases and strengthening the foundation for translational and comorbidity research.

## Results

### Overview of FM-GPT method

Figure 1A shows a schematic overview of our FM-GPT method and Figure 1B shows the directed acyclic graph (DAG) of the full Bayesian model (see Methods for details). Like many other TWAS methods, FM-GPT first trains the genotype-gene prediction models in a reference panel to estimate genotype-gene weight matrix (*γ*), which is then used to impute GReX in the GWAS cohort (Fig 1A-1B (1)). The imputed GReX is subsequently used to assess association with the phenotypes of interest.

**Figure 1.**
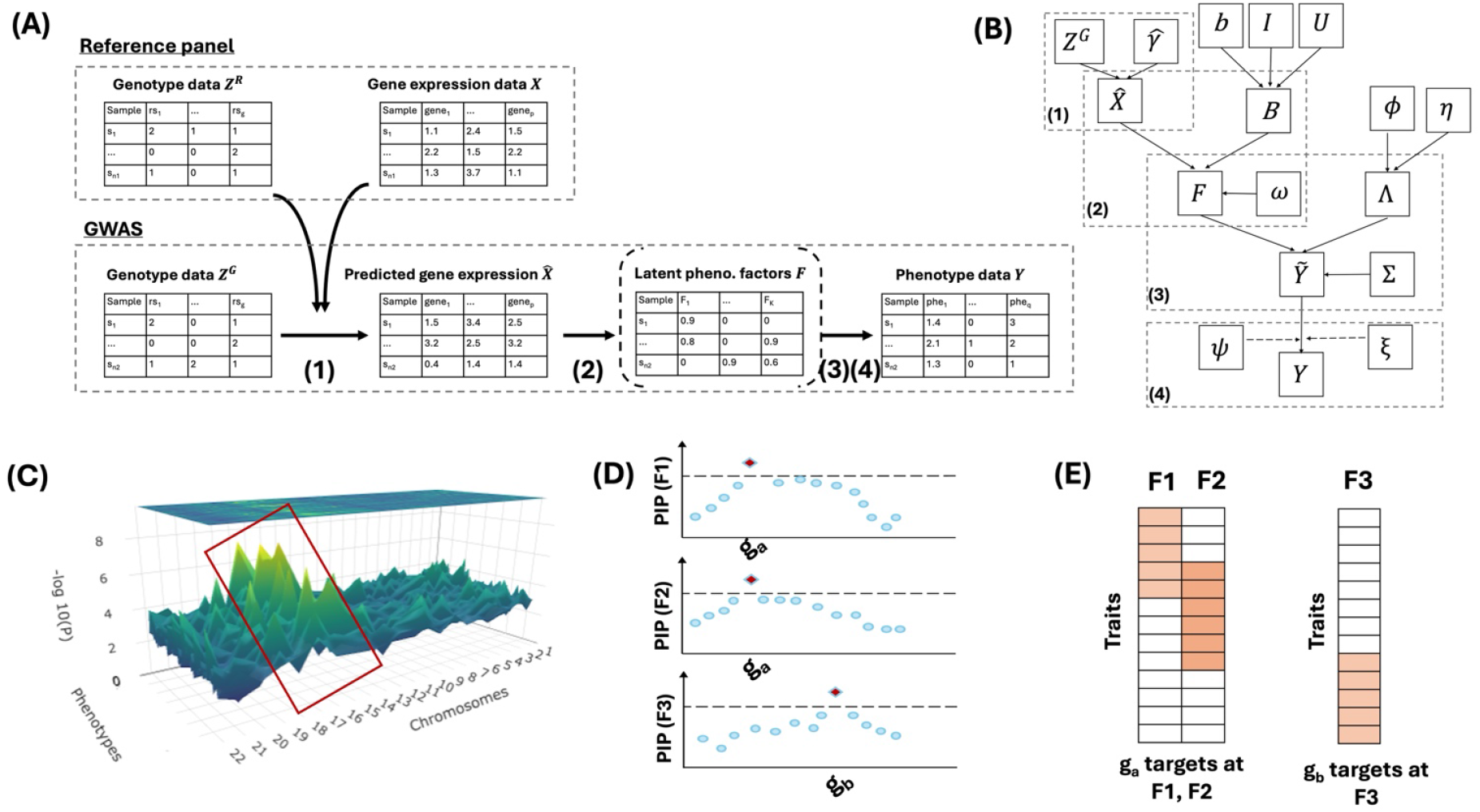
(A) A schematic overview of the FM-GPT method. (B) A directed acyclic graph (DAG) of the FM-GPT method. (C) Demonstration of phenome-wide TWAS results, highlighted regions are where we will apply fine mapping. (D-E) Main outputs from the FM-GPT method: the putatively causal genes for the phenotype factors (D); the loadings of phenotypes on each phenotype factor (E).

FM-GPT performs fine mapping in one genomic region at a time, where the genomic regions are defined based on LD structure (e.g. LD blocks defined by LDetect(20)). For a single phenotype, regions of interest are typically selected from univariate TWAS results based on genome-wide signals. When multiple phenotypes are jointly analyzed, we first obtain the univariate TWAS results over the whole genome for each phenotype and then apply standard meta-analysis approaches, such as Fisher’s method for combining p-values across traits, to identify genomic regions showing evidence of association (e.g. highlighted part in the 3D Manhattan plot in Fig 1C).

To jointly analyze multiple correlated phenotypes, FM-GPT integrates a Bayesian sparse factor analysis model within a Bayesian variable selection framework. The formulation enables supervised dimension reduction of the phenotype space while allowing gene expression signals to guide the identification of latent phenotype factors (Fig 1A-1B (2)(3)). Within each genomic region, the GReX of all candidate genes is modeled simultaneously to prioritize putatively causal genes associated with these latent phenotype factors (i.e. those with highest posterior inclusion probability (PIP) as in Fig 1D). In addition, sparse factor loadings are used to identify the subset of phenotypes most strongly influenced by the genes associated with each latent factor for revealing phenotype-specific gene regulation and improving the interpretability of the inferred factors (Fig 1E). Lastly, FM-GPT accommodates phenotypes of mixed data types - including continuous, binary and count outcomes - through a data augmentation strategy embedded within the full Bayesian model (Fig1A-1B (4)). We leave all technical details of the model, estimation and inference to the Methods section 1-4.

### FM-GPT accurately detects true causal genes across multiple traits while controlling false positives

To assess the performance of our proposed method, we conducted extensive simulations under a variety of scenarios and compared against representative competing methods from several categories. These included TWAS fine mapping methods GIFT (14) and MVIWAS(21), GWAS fine mapping methods PAINTOR(17) and CAVIAR (22), and a general-purpose Bayesian multivariate fine mapping method mvSuSIE (23) (see Table 1 for a comparison of the main characteristics of these methods). Among these methods, FM-GPT is the only method that allows phenome-wide analysis and accommodates phenotypes of mixed data types. In addition to prioritizing putative causal genes, FM-GPT simultaneously selects relevant phenotypes through sparse factor loadings, improving interpretability. Details on how we implemented the competitive methods can be found in the Supplement S2.

**Table 1.**
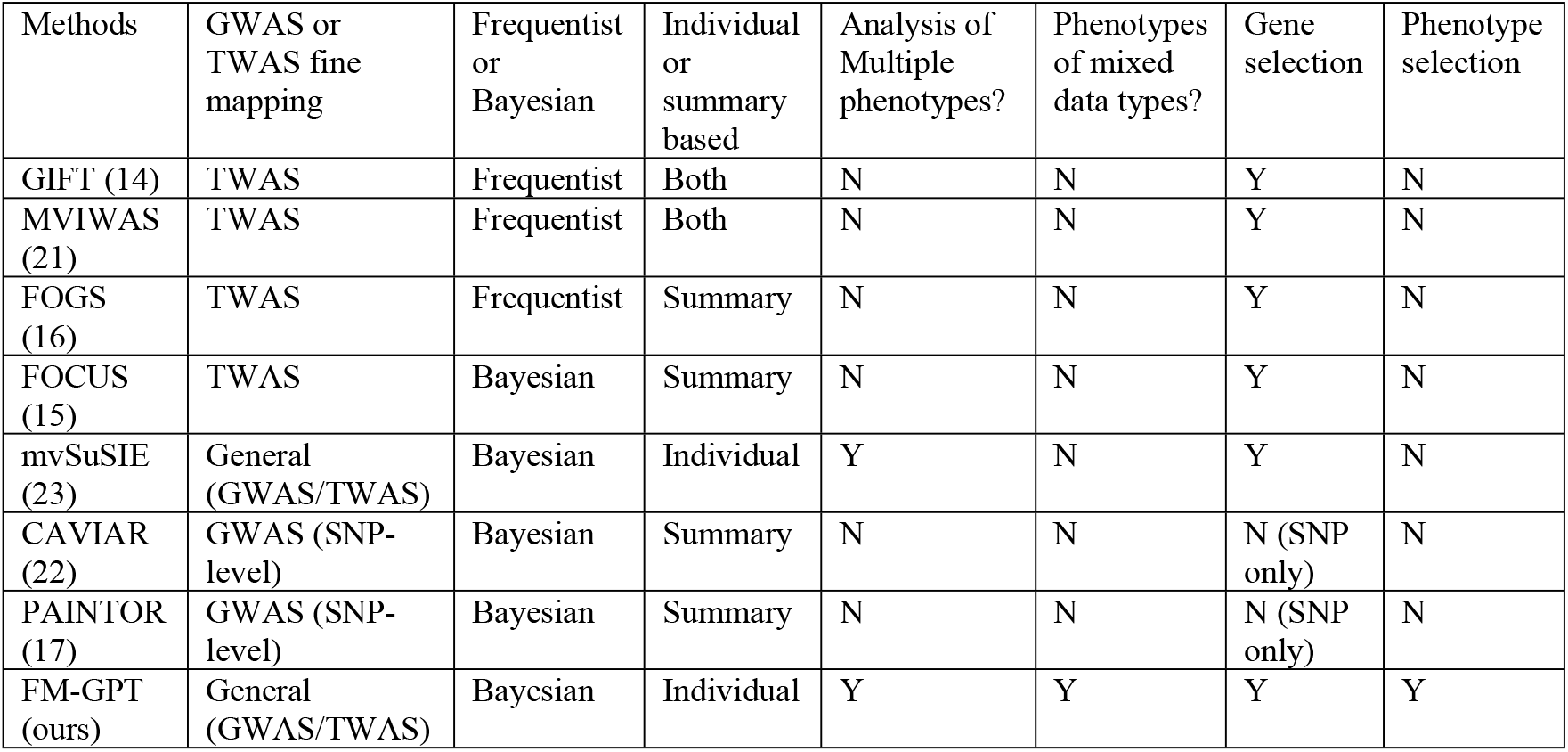
A comparison of main characteristics of representative GWAS/TWAS fine mapping methods/tools.

We considered three simulation settings: (i) homogeneous causality (Scenario 1), where all latent phenotypic factors share the same causal genes; (ii) heterogeneous causality (Scenario 2), where causal genes differ across latent factors; and (iii) a null scenario (Scenario 3) with no true causal genes. Each simulation used a reference panel of size *n*_1_=250, and GWAS sample size *n*_2_=5000, reflecting typical TWAS settings. We simulated 10 genomic regions, each containing 10 genes with an average of 1-2 true causal gene per region. Each gene included 10 cis-SNPs. Phenotypes (q=50) were generated from 1, 3 or 5 latent phenotypic factors and were either continuous or of mixed data types (continuous, binary and count). We further varied phenotypic heritability (i.e. the proportion of phenotypic variance explained by the gene expression) levels (1%, 3%, 5%) for comprehensive assessment. Detailed simulation setup including genotype, gene expression and phenotypic data generation are summarized in the Supplement S1.

Fine mapping problem can be framed as a variable selection problem that aims to identify a parsimonious set of predictors (e.g. SNPs or genes) from a large number of correlated variables(24). Thus, we evaluated the overall variable selection performance using Area Under the ROC Curve (AUC), which is independent of selection thresholds. We also compared the number of true causal genes and the number of false positives identified at a fixed FDR cutoff at 0.1 (FDR-adjusted p-values for the frequentist methods and Bayesian FDR for the Bayesian methods, see details in Methods section 4-5). Because CAVIAR and PAINTOR are designed for SNP-level GWAS fine mapping, their performance was assessed based on identification of cis-SNPs for the causal genes. GIFT and MVIWAS (equivalent to two-stage GIFT, see Methods section 5) are designed for single trait TWAS fine mapping, so we first applied factor analysis to the phenotypes (GIFT with one factor, three factors and five factors) and then performed gene-level analyses for each phenotypic factor, followed by meta-analysis. To mitigate potential bias introduced by factor analysis, in the real-data example 2 using EHR derived phenotypes, we additionally ran GIFT on each of the top phenotypes independently for comparison. mvSuSIE accommodates multivariate phenotypes but assumes continuous outcomes only thus cannot handle mixed data types. Finally, we also evaluated the ability of FM-GPT to recover the correct sparse factor loading structures using F-scores, in comparison with conventional exploratory factor analysis (EFA; low loadings in EFA are suppressed to zeros for fair comparison; see Table S1a-c).

Under homogeneous causality case (Scenario 1), FM-GPT consistently achieved the highest AUC across varying numbers of latent factors and heritability levels, for both continuous and mixed-type phenotypes (Fig 2A-B), while maintaining well-controlled false positives (Fig 2C-D). GIFT and MVIWAS showed high sensitivity but at the cost of substantially inflated false positives. PAINTOR and CAVIAR, which were designed for GWAS fine mapping, were generally underpowered. Although mvSuSIE effectively controlled false positives, its overall variable selection performance was inferior to FM-GPT, particularly when the number of latent factors was small or signal strength was low (i.e., low heritability). In the heterogeneous setting (Scenario 2), performance declined for all methods; however, FM-GPT remained the top-performing approach and demonstrated more pronounced advantages across all conditions (Fig 3A-D). Beyond gene prioritization, FM-GPT accurately recovered the subset of phenotypes with nonzero loadings, outperforming standard EFA (Table S1a-c). Under the null scenario (Scenario 3), only FM-GPT, mvSuSIE and CAVIAR adequately controlled false positive rates, whereas other methods exhibited substantial inflation (Fig S1). In light of the simulation results, we primarily compare FM-GPT to GIFT, MVIWAS and mvSuSIE for real data examples.

**Figure 2.**
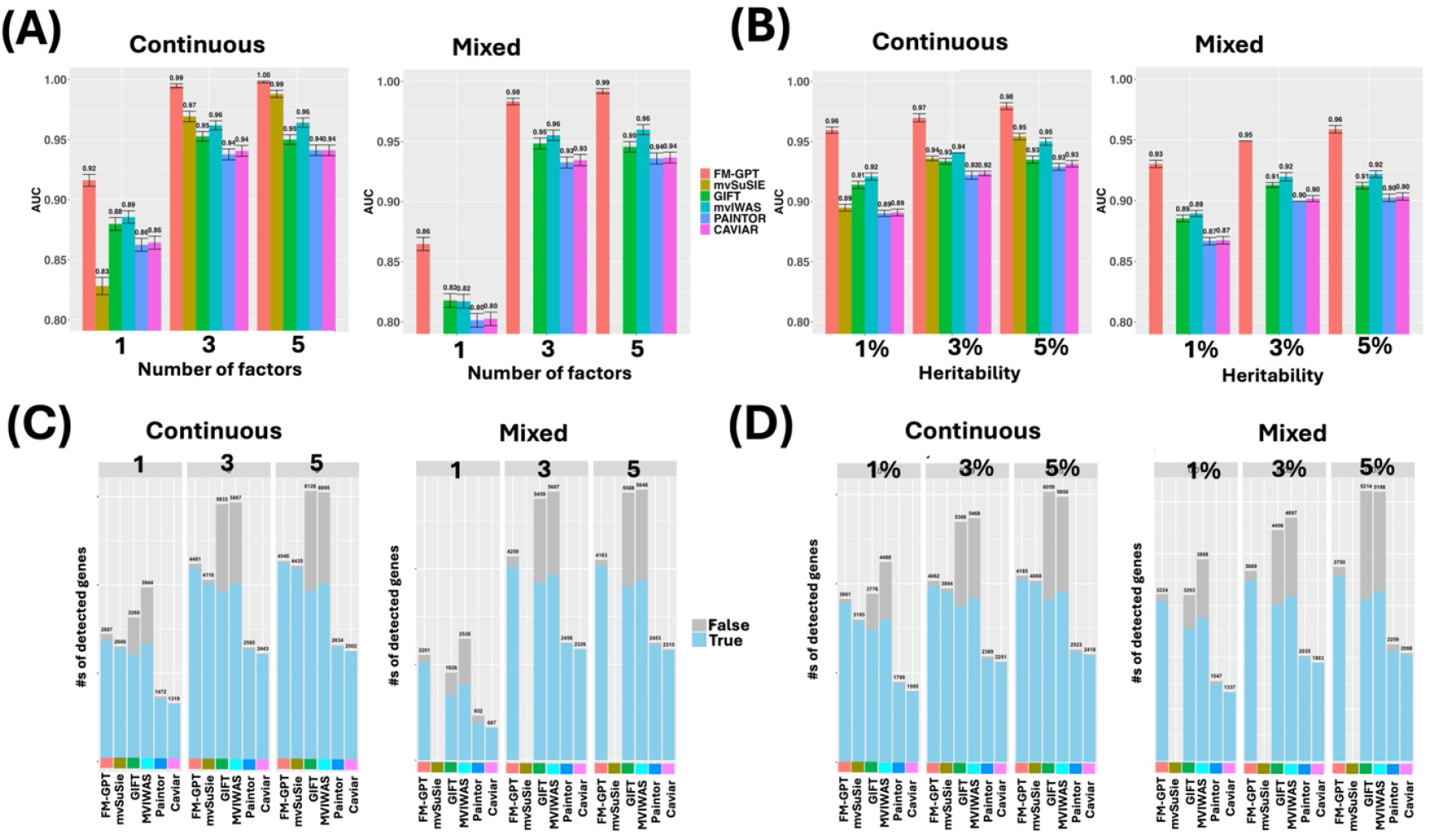
Simulation results for scenario 1 with homogeneous causality. (A) AUCs of all methods with varying number of factors for both continuous phenotypes only and phenotypes of mixed data types. (B) AUCs of all methods with varying heritability levels for both continuous phenotypes only and phenotypes of mixed data types. (C) Number of true positives vs false positives detected by all methods at same FDR cutoff (0.1) with varying number of factors for both continuous phenotypes only and phenotypes of mixed data types. (D) Number of true positives vs false positives detected by all methods at same FDR cutoff (0.1) with varying heritability levels for both continuous phenotypes only and phenotypes of mixed data types. *mvSuSIE is removed from mixed outcome plots as they cannot handle mixed data types.

**Figure 3.**
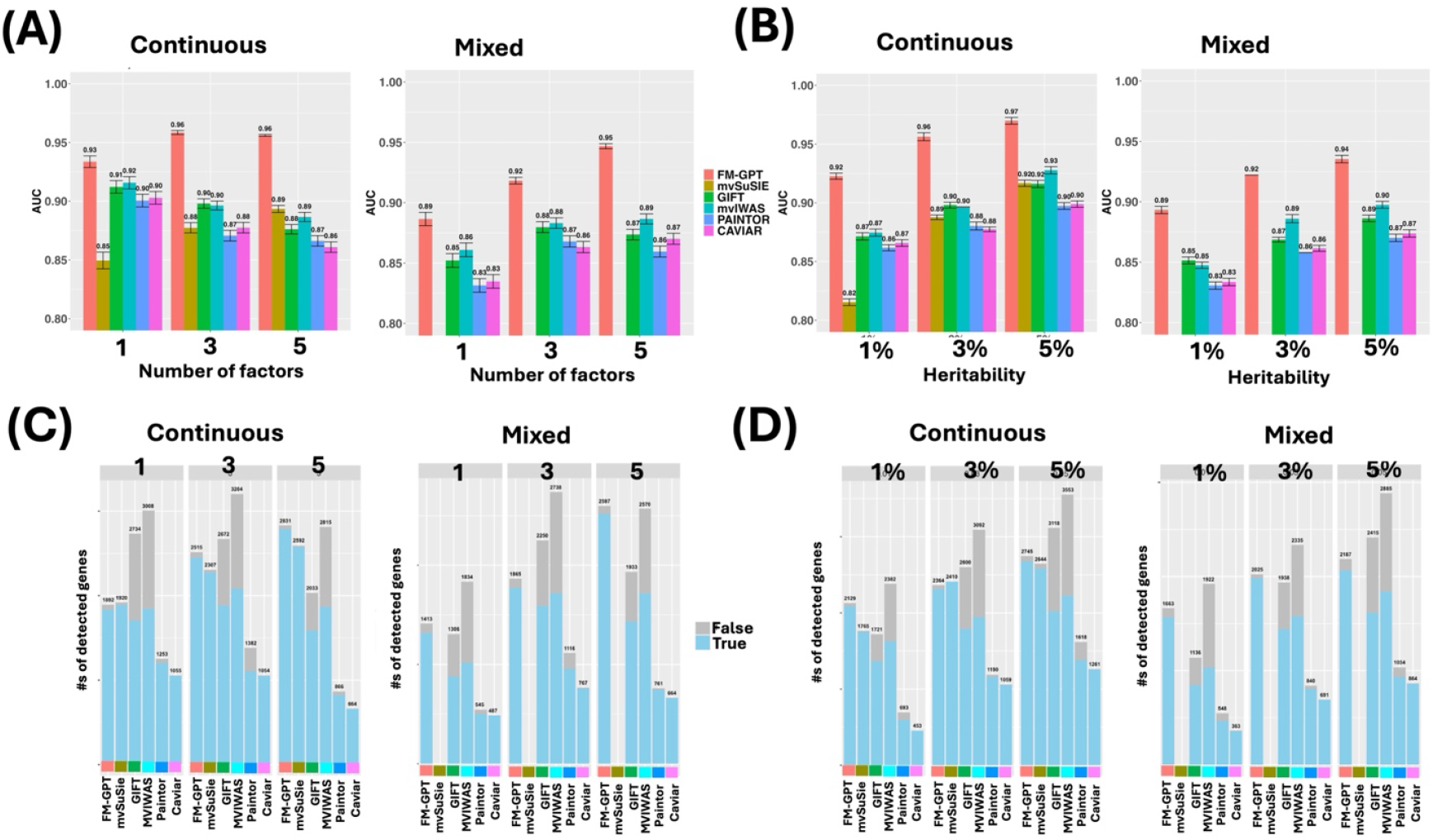
Simulation results for scenario 2 with heterogeneous causality. (A) AUCs of all methods with varying number of factors for both continuous phenotypes only and phenotypes of mixed data types. (B) AUCs of all methods with varying heritability levels for both continuous phenotypes only and phenotypes of mixed data types. (C) Number of true positives vs false positives detected by all methods at same FDR cutoff (0.1) with varying number of factors for both continuous phenotypes only and phenotypes of mixed data types. (D) Number of true positives vs false positives detected by all methods at same FDR cutoff (0.1) with varying heritability levels for both continuous phenotypes only and phenotypes of mixed data types. *mvSuSIE is removed from mixed outcome plots as they cannot handle mixed data types.

### FM-GPT identifies genes with pleiotropic effects on brain-wide cortical thickness measures from structural MRI data in UKB

We applied FM-GPT to two real data applications in UK Biobank (UKB). In the first example, we investigated the genetic architecture of brain structure using structural MRI data from the UKB. Cortical thickness (CT), defined as the distance between the gray and white matter boundaries, is a key neuroanatomical phenotype linked to cognitive function, neurodevelopment, and a range of neurological and psychiatric disorders(25-27). CT is also highly heritable, with both global and region-specific genetic influences(28, 29). Here, we aimed to leverage FM-GPT to perform TWAS and fine mapping analyses of regional CT across the entire cerebral cortex, with the goal of prioritizing causal genes underlying brain-wide variations in cortical structure.

We analyzed CT measures of 66 cortical regions (see Table S2 for a list of these brain regions) defined by the Desikan-Killiany Atlas(30), derived using FreeSurfer(31). After restricting to genetically unrelated individuals with complete genetic and covariate data (age, sex, body mass index (BMI) and top 10 genetic principal components), the final sample included n_2_=26,124 individuals. As expected, regional CT measures were highly correlated, reflecting substantial shared variation across the cortex (Fig 4A).

**Fig 4.**
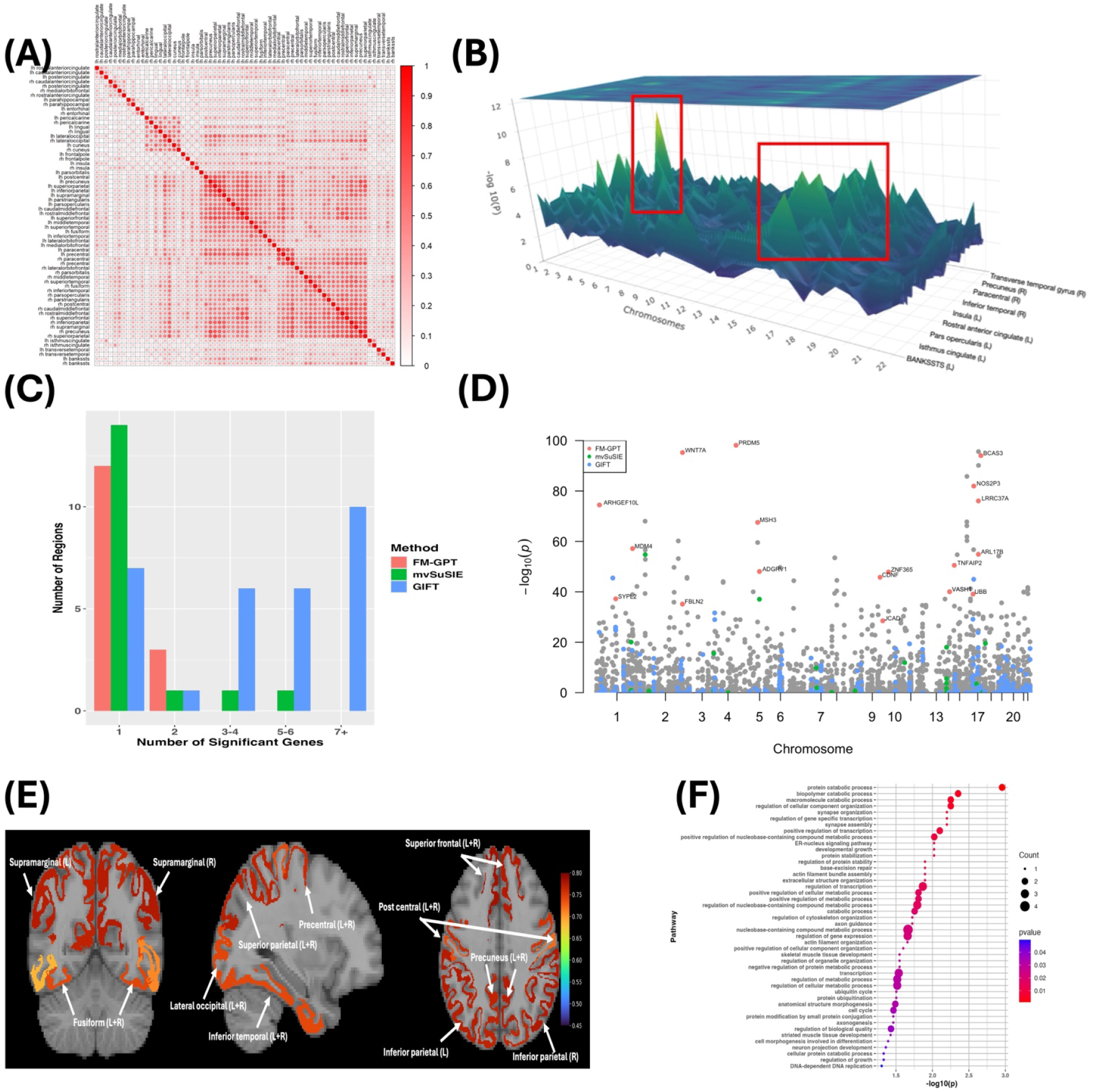
TWAS and fine mapping results for genetic study of 66 regional cortical thickness (CT) measures from UKB. (A) The Phenotypic correlation of the 66 regional CT measures; (B) 3D Manhattan plot of TWAS p-values (most significant genomic regions highlighted); (C) Proportion of regions that harbor different number of causal genes by different methods; (D) Manhattan plot of meta-TWAS Fisher’s method p-values and the putative causal genes identified by each fine mapping method; (E) Brain regions with the largest factor loadings; (F) Pathway enrichment analysis on FM-GPT selected genes results (p<0.05).

We first conducted univariate TWAS using 13 brain tissues from GTEx(1), for each of the 66 regions (see Fig 4B for the 3D Manhattan plot; Table S3) adjusting for covariates including age, sex, BMI and the top 10 principal components. We then aggregated evidence across regions using Fisher’s method and selected 760 genes (p<1e-8) for downstream analysis. Using LDetect(20), we selected 355 genomic regions containing at least one TWAS significant gene as regions of interest which we would fine map next (Table S4).

FM-GPT identified 18 putative causal genes across 16 genomic regions at a Bayesian false discovery rate (BFDR) of 0.15 (Table 2). In comparison, mvSuSIE identified 25 genes, GIFT and MVIWAS (assuming a single factor) identified 164 and 174 genes, respectively at the same FDR cutoff (two genes selected by all methods; Table S4). Notably, FM-GPT demonstrated substantially greater specificity, with a higher proportion of regions containing only 1-2 prioritized genes and greatly narrowed down the set size of putatively causal genes by 28-90% compared with other methods (Fig 4C, Table 2). Moreover, genes prioritized by FM-GPT were supported by stronger TWAS signals overall, whereas competing methods frequently selected genes with weak or non-significant associations, suggesting inflated false positives (Fig 4D).

**Table 2.** Summary of fine mapping results for the 355 regions by different methods in the cortical thickness example.

| Methods | Number of significant genes from fine mapping (FDR<0.15) | Number of regions with 1 causal gene | Number of regions with 2 causal genes | Number of regions with 3-4 causal gene | Number of regions with 5-6 causal gene | Number of regions with 7+ causal gene |
| --- | --- | --- | --- | --- | --- | --- |
| FM-GPT | 18 | 12 | 3 | 0 | 0 | 0 |
| mvSuSIE | 25 | 14 | 1 | 1 | 1 | 0 |
| GIFT | 164 | 7 | 1 | 6 | 6 | 10 |
| MVIWAS | 174 | 45 | 15 | 14 | 1 | 4 |

Among them, FM-GPT identified critical pleiotropic genes localized to a region on chromosome 17, including *BCAS3, LRRC37A, NOS2P3, ARL17B* and *UBB*, participated in neuronal morphology and cortical organization, highlighting a coordinated genetic program influencing distributed cortical architecture (32, 33). Importantly, FM-GPT revealed that these genes act through a shared latent factor encompassing 46 cortical regions (Table S5, see the regions with highest loadings in Fig 4E), indicating a common genetic program influencing widespread cortical architecture.

This finding highlights a key advantage of FM-GPT: rather than analyzing each region independently or relying on global summary measures, our approach jointly models brain-wide phenotypes and directly identifies shared causal genes underlying coordinated variation across regions. In contrast to prior studies that examined global mean CT or region-specific GWAS separately(32, 33), FM-GPT provides a unified and interpretable framework for uncovering the genetic basis of distributed brain structure. Finally, pathway analysis of the prioritized genes using Gene Ontology Biological Process database (34) revealed significant enrichment in biological processes related to protein catabolic process, protein ubiquitination and synaptic function (Fig. 4F; Table S6), underscoring the role of neuronal maintenance and synaptic integrity in shaping cortical morphology.

### FM-GPT identifies genes that influence multiple medial conditions derived from EHR data

In a second application, we applied FM-GPT to fine-map genetic associations across a broad spectrum of disease phenotypes derived from EHR data in UKB. EHR data provide a powerful resource for large-scale phenome-wide analyses but present substantial analytical challenges due to their high dimensionality, heterogeneous data types (e.g., binary diagnoses, counts, continuous traits), and complex correlation structure across diseases. These features limit the applicability of conventional fine-mapping approaches, which typically assume homogeneous phenotype types and analyze each trait independently.

We extracted 19,190 ICD10-based disease codes and performed systematic preprocessing to derive 1,403 phenotypes of mixed data types (primarily binary and count), grouped into 16 major disease categories (e.g., circulatory, metabolic, digestive, and respiratory systems)(35). To ensure adequate statistical power and avoid extreme case–control imbalance, we retained 169 phenotypes with prevalence >1% among n_2_=245,687 individuals for downstream TWAS and fine-mapping analyses (Table S7; see detailed steps on processing these EHR-derived phenotypes in Supplement S3).

We first conducted univariate TWAS for each phenotype using all 50 tissues from GTEx(1), adjusting for key covariates, e.g. age, sex, BMI (Fig 5A; Table S8). We then focused on 39 phenotypes with at least one gene showing genome-wide significant association (p < 1e-6), spanning major disease domains including circulatory, digestive, metabolic, respiratory and genitourinary systems. These phenotypes exhibited substantial correlation both within and across clinical categories (Fig 5B), motivating a joint modeling framework. We aggregated TWAS signals across phenotypes using Fisher’s method and defined 297 genomic regions containing at least one significant gene (Table S9) for fine mapping.

**Fig 5.**
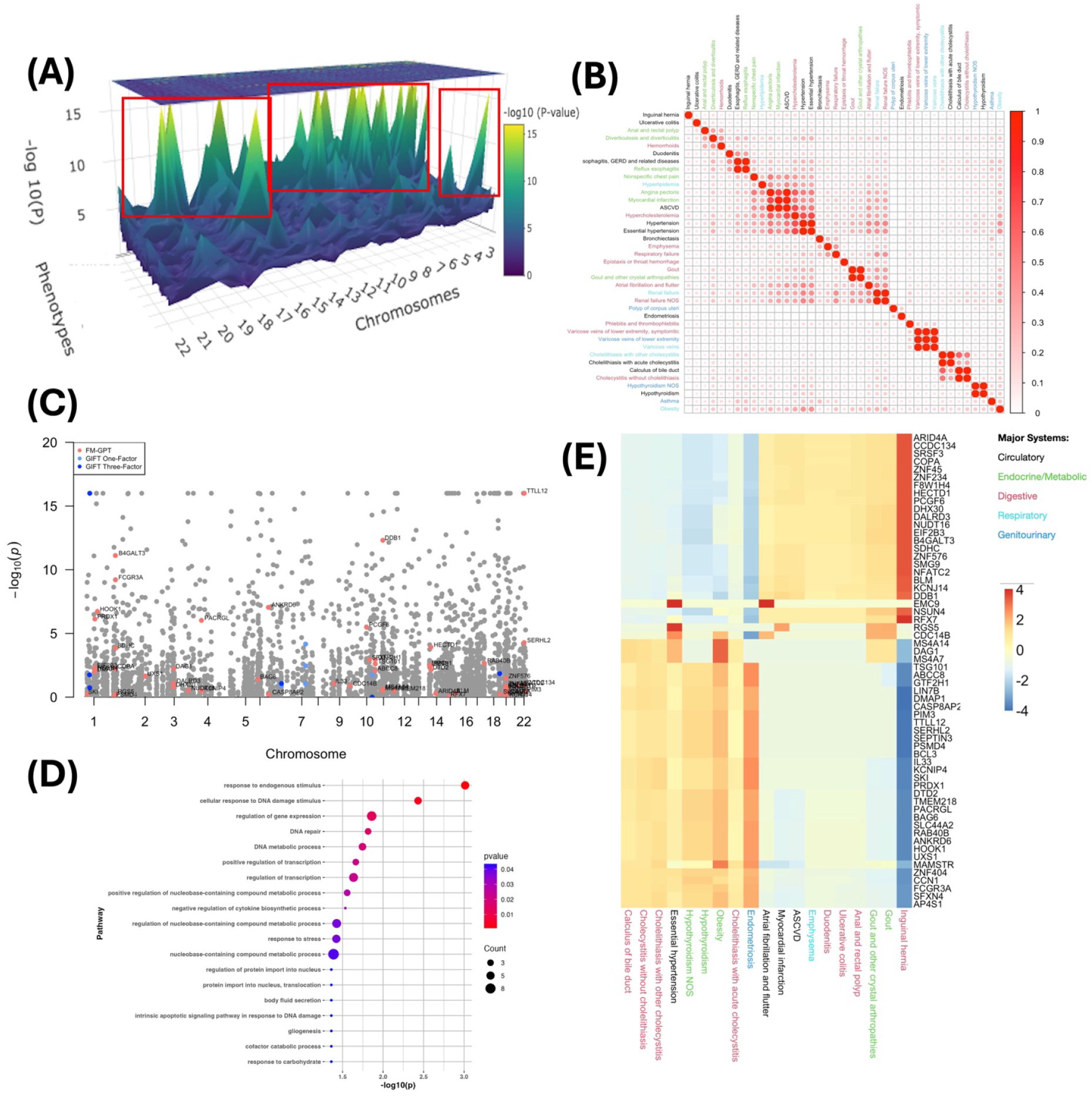
TWAS and fine mapping results for genetic study of EHR-derived phenotypes from UKB. (A) 3D Manhattan plot of TWAS p-values (most significant genomic regions highlighted); (B) The Phenotypic correlation of the 39 EHR-derived medical conditions; (C) Manhattan plot of meta-TWAS Fisher’s p-values and the putative causal genes identified by each fine mapping method; (D) Pathway enrichment analysis on FM-GPT selected genes results (p<0.05); (E) Heatmap of identified putative causal genes and the loadings of phenotypes they target at.

FM-GPT identified 60 putative causal genes at a BFDR of 0.15 (Fig 5C). Since the phenotypes are of mixed data types (primarily binary and count) in this example, mvSuSIE failed to run. GIFT and MVIWAS with one phenotypic factor or three phenotypic factors (see Methods section 5), as well as univariate GIFT independently on top 10 phenotypes ranked by TWAS signals were applied for comparison. As compared to FM-GPT, GIFT and MVIWAS tend to select genes with weak or non-significant TWAS associations (Fig 5C, Table S9). GIFT on univariate phenotype has severe inflation of false positives with hundreds to thousands of genes detected as putative causal, which undermines the reliability and interpretability of fine-mapping results (Table S9). Notably, our method revealed both shared genetic effects across phenotypes and phenotype-specific contributions, enabled by its joint modeling and sparse factor structure. The inferred latent factors captured structured heterogeneity across disease domains, with two major axes of variation emerging (Fig 5E, Table S10). One axis was driven by cardiovascular and inflammatory conditions, including atrial fibrillation, myocardial infarction, and ulcerative colitis, whereas the opposing axis was enriched for metabolic and hepatobiliary phenotypes, such as gallstone-related disorders, hypothyroidism, and obesity. Importantly, genes loading onto these factors exhibited distinct functional profiles. Genes associated with the cardiovascular-inflammatory axis were enriched for roles in transcriptional regulation, RNA processing, and genome maintenance (e.g., *ARID4A, SRSF3, BLM, DDB1*)(36, 37), whereas those associated with the metabolic-hepatobiliary axis included key immune and inflammatory regulators (e.g., *IL33, FCGR3A, BCL3*)(38, 39) (Fig. 5D, Table S11). These results suggest a pleiotropic genetic architecture in which shared regulatory pathways influence multiple disease domains, while also allowing for directionally distinct effects across phenotypic groups.

This pattern is suggestive of a potential immune-metabolic trade-off(40), whereby genetic regulation may coordinate energy allocation between immune function and metabolic processes, however, this remains hypothesis-generating as the directionality of latent factor loadings could be inherently arbitrary. More broadly, these results illustrate how coordinated yet heterogeneous genetic effects across disease domains can be captured through joint modeling. Such patterns are difficult to detect using conventional single-trait or univariate fine-mapping approaches. By simultaneously capturing shared and phenotype-specific genetic influences across diverse and mixed-type traits, FM-GPT provides a flexible and interpretable framework for investigating the genetic architecture of complex, multi-system diseases.

## Discussion

In this study, we developed FM-GPT, a Bayesian fine-mapping framework for phenome-wide TWAS that jointly prioritizes putatively causal genes and latent phenotype factors when multiple correlated outcomes are analyzed together. By combining Bayesian variable selection with sparse supervised factor analysis, FM-GPT is designed to recover both shared and phenotype-specific patterns of genetic regulation across traits while retaining interpretability at the phenotype level. Our simulation studies demonstrate that leveraging phenome-wide information in a joint model can accurately recover the true causal genes without inflating the false positives. In UKB applications, FM-GPT greatly narrowed down the set size of putatively causal genes compared to other methods, and highlighted genes with pleiotropic effects on regional cortical thickness measures across the cerebral cortex, as well as genes that influence multiple EHR-derived medical conditions spanning circulatory, metabolic, digestive, respiratory and genitourinary systems, and reveal potential immune-metabolic gene regulatory trade-off. These results demonstrate that FM-GPT enables the discovery of shared genetic architecture across the phenome, uncovering both universal and phenotype-specific mechanisms that are not apparent from single-trait analyses or conventional fine-mapping approaches.

Existing fine-mapping approaches are largely designed for single-phenotype analyses, limiting their direct applicability to phenome-wide settings. Applying these methods independently across traits introduces a substantial multiple testing burden and often leads to reduced power, particularly when phenotypes are correlated(41, 42). Conversely, dimension-reduction approaches such as factor analysis or principal component analysis applied a priori may oversimplify heterogeneous trait architectures, and the resulting latent factors are not guaranteed to align with underlying genetic effects(43). FM-GPT bridges these two extremes by jointly modeling latent factor structure and gene selection within a unified framework. By allowing genetic signals to inform factor construction, the method identifies latent phenotypic structures that are more directly linked to underlying biology. This joint modeling strategy improves power to detect shared genetic effects while preserving heterogeneity across traits. In addition, FM-GPT accommodates mixed data types and enforces sparse factor loadings, enhancing both interpretability and scalability for complex phenome-wide analyses.

The FM-GPT framework can be extended in several important directions. Firstly, incorporating multi-omics QTL resources beyond expression QTLs used in TWAS, such as splicing QTLs, methylation QTLs, and chromatin accessibility QTLs, could substantially improve resolution in prioritizing regulatory mechanisms and causal genes(44-46). Secondly, though our current implementation integrates expression prediction models across multiple tissues to maximize statistical power, future extensions could explicitly incorporate tissue- and cell-type-specific regulatory architectures. Such approaches may enhance biological interpretability, particularly for traits with well-defined cellular contexts, as demonstrated in recent single-cell and tissue-resolved genetic studies(47, 48). Thirdly, the current model does not explicitly account for causal relationships among phenotypes. Extending the framework to jointly model genetic effects and directed dependencies between traits, potentially through integration with causal inference or structural equation modeling approaches, could provide deeper insight into mechanisms of pleiotropy and mediation(43, 49, 50). Lastly, further methodological development is warranted to scale to increasingly large phenomic datasets. In particular, extending FM-GPT to operate directly on GWAS summary statistics would broaden its applicability, while variational Bayesian approximations could offer additional gains in computational efficiency relative to Gibbs sampling(51). For settings involving very large numbers of genes, incorporating principled feature-screening or preselection strategies may further improve scalability(52). As phenomic and multi-omics resources continue to expand, methods that jointly model genetic effects across diverse traits will become increasingly critical. By integrating information across traits while preserving heterogeneity, FM-GPT provides a flexible and extensible framework that will play an increasingly important role in translating association signals into biological meaningful insights. The R package to implement FM-GPT method is freely available at www.github.com/tacanida/fm-gpt.

## Methods

In the following sections, we present an overview of the FM-GPT method, including the model formulation, prior specification and the data augmentation strategy for handling mixed data types, as well as the procedure for selecting putative causal genes. We also describe the data and the cohort used in the two application examples.

### 1. Model of FM-GPT

Consider a reference eQTL dataset (e.g. GTEx) with *n*_1_ individuals and a GWAS dataset with *n*_2_ individuals and *q* measured phenotypes. Following standard convention in fine mapping(24), we first identify genomic regions of interest based on phenome-wide TWAS results (e.g. via meta-analysis of univariate TWAS p-values). Suppose there are *T* regions of interest identified, and let *p*_*t*_ denote the number of genes in *t*-th region (1 ≤ *t* ≤ *T*). We consider genes within each genomic region (based on LD blocks defined by LDetect(20)) as potentially predictors of the phenotypes under study, analyzing one region at a time. Without loss of generality, we will introduce the model formulation (section 1), prior specification (section 2) and data augmentation (section 3) using *t*-th genomic region as an example, and the same setting applies to all *T* regions of interest. For selecting putative causal genes using Bayesian false discovery rate (section 4), we will consider all genes and all genomic regions.

Let *x*_*j*_ denote the *n*_1_-vector of gene expression levels for the *j*-th gene in the reference dataset, for *j* = 1,2,…,*p*_*t*_. Let 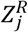 denote the *n*_1_x *g*_*j*_ genotype matrix of the *g*_*j*_ cis-SNPs for the jth gene in the reference dataset and 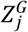 denote the corresponding *n*_2_x *g*_*j*_ genotype matrix in the GWAS dataset. Let *γ*_*j*_ denote the *g*_*j*_x 1 vector of cis-SNP effect sizes (i.e. the weight vector) for the jth gene and let 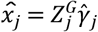 the corresponding predicted GReX in the GWAS dataset for *j* = 1,2,…,*p*_*t*_.

Let *Y* = {*y*_*ik*_;*i* = 1,2,…*n*_2_,*k* = 1,2,…,*q*} denote the *n*_2_x *q* phenotype matrix in the GWAS dataset, the *q* phenotypes are typically correlated and are assumed to be represented by *m* latent factors. Let *B* = {*β*_*jl*_ ;*j* = 1,2,…*p*_*t*_,*l* = 1,2,…,*m*} denote the *p*_*t*_x *m* matrix of gene effects on phenotype factors. Let *F* denote the *n*_2_x *m* matrix of factor scores, where each column *f*_*l*_ corresponds to the lth factor (*l* = 1,2,…,*m*), and let Λ denote the *q*x *m* factor loading matrix, where each column *λ*_*l*_ represents the loadings for the lth factor. The phenotypes may be of mixed data types, such that each column of *Y* can be continuous or discrete. We assume that each column of 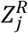 and 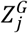 has been standardized to have mean zero and unit variance.

Figure 1B shows a DAG of the full joint model. The joint model considers the following equations for each *t*-th (1 ≤ *t* ≤ *T*) genomic region (the same model applies over all *T* regions):

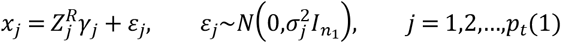

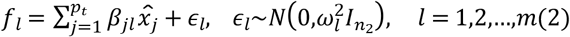

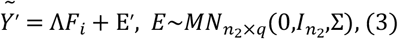

where each element in 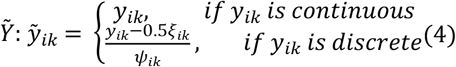, for *i* = 1,2,…,*n*_2_; *k* = 1,2,…,*q*. 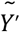 and E′ denote the transpose of 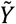 and E, respectively, *F*_*i*_ is the *i*-th row of *F* (*i* = 1,2,…,*n*_2_).

Equation (1) specifies a per-gene regression model describing the relationship between the expression of the jth gene and its cis-SNP regulators in the reference panel dataset (i.e. the training stage in TWAS method(3)). ε_*j*_ denotes the error that follows 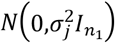, where 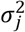 is a gene specific variance. Because the number of cis-SNPs *g*_*j*_ is typically larger than the sample size *n*_1_, sparsity must be imposed on *γ*_*j*_ to ensure model identifiability. Here, we adopt a Bayesian sparse linear model with a spike-and-slab prior on *γ*_*j*_ to select nonzero effects (see section 2 for prior specification). When multiple tissues are available in the training data, equation (1) can be extended by incorporating a multivariate spike-and-slab prior to jointly model SNP effects across tissues, thereby improving statistical power(9). The training stage can be computationally intensive when many SNPs map to each gene, and access to raw reference panel data may be limited. As an alternative, users can directly leverage pre-trained weight matrices from existing models (e.g. PrediXcan(5), MultiXcan(8) or UTMOST(9)) to impute GReX in the GWAS dataset.

Equation (2) specifies a per-factor regression model describing the relationship between GReX for each gene and the latent phenotype factors in the GWAS dataset. The fine-mapping problem can be framed as a variable selection problem that aims to identify a parsimonious set of predictors (e.g., SNPs or genes) from a large number of correlated variables. In our setting, the goal is to identify putative causal genes that influence multiple phenotypes among all genes located within a genomic region of interest. *ϵ*_*l*_ denotes the error term assumed to be independently and identically distributed as 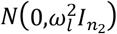, where 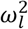 is the variance of the lth factor. *β*_*jl*_ is the parameter of interest in fine mapping, as it characterizes gene effects on phenotypes and enables the identification of true causal genes from a large set of candidates, including those with null effects. We consider both factor-specific and omnibus hypotheses. Specifically, for each gene-factor pair, we test *H*_0_:*β*_*jl*_ = 0 vs. *H*_*a*_:*β*_*jl*_ ≠ 0 for 1 ≤ *j* ≤ *p* and 1 ≤ *l* ≤ *m*. We also consider an omnibus test of gene effects over all factors, defined as *H*_0_:⋂_*l*_{*β*_*jl*_ = 0} vs. *H*_*a*_:⋃_*l*_{*β*_*jl*_ ≠ 0} for 1 ≤ *j* ≤ *p*. As our method is formulated within a full Bayesian hierarchical framework, we use posterior inclusion probabilities and the corresponding Bayesian false discovery rate to prioritize putative causal genes(53) (see Section 4).

Equation (3) specifies a factor model that describes the relationship between a large number of observed phenotypes and a smaller number of latent phenotype factors. 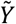 denotes a transformed version of the observed phenotype matrix *Y*, used to accommodate both continuous and discrete traits as described in equation (4). We impose sparsity-inducing priors on Λ to obtain sparse factor loadings (see Section 3). As in standard latent factor models, we assume that the error matrix 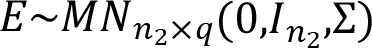, where Σ is assumed to be a diagonal matrix *diag*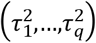 with 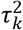 representing the phenotype-specific variance for the *k*-th phenotype. The parameters *ξ*_*ik*_ and *ψ*_*ik*_ arise from the data augmentation procedure described in Section 3 data augmentation strategy. Unlike standard unsupervised factor analysis which assumes a single global set of latent factors, our model allows the latent factors to be specific to each genomic region under study. Consequently, phenotypes are projected into a gene-dependent latent space (i.e., a supervised factor model), in which the factor loadings are influenced by the genetic effects of causal genes within that region. This formulation enables us to characterize how different genes influence multiple phenotypes (e.g., Fig. 5E), providing insight into potentially coherent biological mechanisms and facilitating hypothesis generation.

Compared with multi-stage modeling approaches, our joint model offers several advantages. First, it enables joint modeling of multiple genes and multiple phenotypes, along with multi-level selection at the phenotype, factor, and gene levels, which facilitates the identification of complex and potentially heterogeneous regulatory patterns between genes and phenotypes. Secondly, unlike commonly used unsupervised factor models—where latent factors are derived independently of predictors—our approach employs a supervised factor model in which the mapping from phenotypes to latent factors is partially driven by putative causal genes. In addition, sparsity in the factor loadings promotes the selection of relevant phenotypes and improves the interpretability of the latent factors. Third, our joint model can accommodate mixed types of phenotypes, addressing a critical challenge in genetic studies involving multiple traits(24).

The primary parameters of interest in the model are Λ and *B*. The matrix *B* is used to identify which genes are inferred to be causal and to determine the factors they influence. The loading matrix Λ identifies the subset of the phenotypes of that load onto each latent factor.

### 2. Prior Setting

For the full model, we consider three sparsity-inducing priors to encourage selection of cis-SNP effects on gene expression 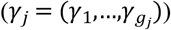 in the reference dataset, selection of gene effects on latent factors (*β*_*jl*_), and sparse loadings of observed phenotypes on each latent factor (*λ*_*l*_).

For each element in *γ*_*j*_, we propose a standard spike-and-slab prior for the effect of *d*-th SNP on gene expression, *γ*_*d*_, for *d* = 1,2,…,*g*_*j*_, as in many Bayesian TWAS models(3, 13):

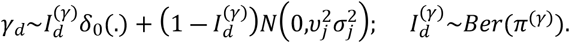

*δ*_0_(.) is the point mass at zero. The spike-and-slab prior enables exact variable selection by shrinking small effect sizes to zero through a mixture of a point mass at zero (spike) and a continuous distribution over the real line (slab). For the hyperparameters, we assume standard uniform and Jeffrey’s priors: 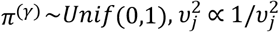

For *β*_*jl*_, we specify the indicator variable priors (equivalent to spike-and-slab priors(54, 55)) to select putative causal genes for each phenotype factor:

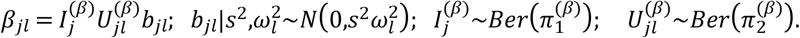

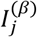 and 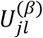 are two indicator variables serving distinct roles in feature selection: 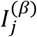 controls gene-level selection and 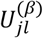 determines whether a gene is active for a specific factor. *b*_*jl*_ represents the underlying effect of the *j*-th gene on the lth factor. For the hyperparameters, we assume standard uniform and Jeffrey’s priors: 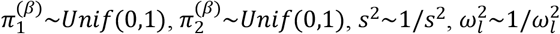.

For *λ*_*l*_, we apply a gamma process shrinkage prior(56) to induce sparsity in the factor loadings, while allowing the number of factors to go to infinity. For *k* = 1,2,…,*q* and *l* = 1,2,…,*m*, we assume the following prior on each loading coefficient:

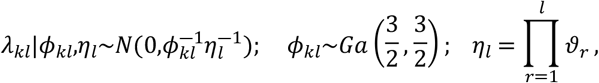

where Ga refers to gamma distribution, *ϑ*_1_∼*Ga*(*a*_1_,1) and *ϑ*_*h*_∼*Ga*(*a*_2_,1) for *h* ≥ 2. In our prior setting, we assume *a*_1_ > 1 and *a*_2_>1, which induces stochastic shrinkage in η_*l*_ and encourages progressively stronger shrinkage for higher-indexed factors. The parameters *ϕ*_*kl*_ and η_*l*_ together define a global-local shrinkage structure on the loadings, preventing any individual loading vector or element from becoming excessively large. Finally, we allow the data to determine the effective number of factors by incorporating an adaptive step within the Gibbs sampler (see Supplement S4 Gibbs sampler).

### 3. Data augmentation strategy for mixed data types

To accommodate phenotypes of mixed types (e.g., continuous, binary, and count), we introduce a data augmentation strategy(57, 58). For *i* = 1,2,…,*n*_2_; *k* = 1,2,…,*q*, we define the transformed variable *y*_*ik*_ as:

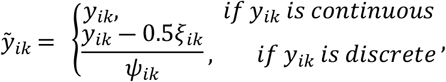

where *ξ*_*ik*_ =1 when *y*_*ik*_ is binary and *ξ*_*ik*_ = *y*_*ik*_ + *r*_*k*_ when *y*_*ik*_ is count data, *r*_*k*_ is the parameter that describes the number of successes in the negative binomial distribution and is updated within the Gibbs sampler. For categorical variables, we either transform them into multiple dummy variables or follow (57) to derive their augmented representations directly. The auxiliary variable *ψ*_*ik*_ is drawn from a Pólya-Gamma (58)distribution *PG*(*ξ*_*ik*_,0) when *y*_*ik*_ is discrete. This data augmentation strategy unifies the likelihood for both continuous and discrete phenotypes, enabling the use of conjugate conditional posteriors within a Gibbs sampling framework.

All the parameters in the full model have closed-form posterior distributions, so we use Gibbs sampling algorithm to update all the parameters (see Supplement S4 Gibbs sampler).

### 4. Selection of the putative causal genes using Bayesian false discovery rate

We will follow the convention in Bayesian fine mapping literature (24) to summarize the posterior distribution of the effect sizes *β*_*jl*_ using marginal posterior inclusion probability (PIP) of each gene. Suppose we test a total of 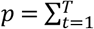 genes across *T* genomic regions. For factor-specific inference, we test *H*_0_:*β*_*jl*_ = 0 vs. *H*_*a*_:*β*_*jl*_ ≠ 0, and define the PIP for the *j*-th (1 ≤ *j* ≤ *p*) gene and the lth factor (1 ≤ *l* ≤ *m*) as *PIP*_*jl*_: = *P*_*jl*_(*H*_*a*_|*data*) = *P*(*β*_*jl*_ ≠ 0|*data*). To account for multiple testing across genes, we define the factor-specific Bayesian false discovery rate (BFDR) for factor *l*: 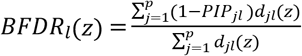, where *d*_*jl*_(*z*) = *I*{*PIP*_*jl*_ < *z*}. We select putative causal genes by controlling *BFDR*_*l*_ at a pre-specified level *α*.

For the omnibus test of gene effects across all factors, *H*_0_:⋂_*l*_{*β*_*jl*_ = 0} vs. *H*_*a*_:⋃_*l*_{*β*_*jl*_ ≠ 0}, we define the gene-level PIP as *PIP*_*j*_: = *P*_*jl*_(*H*_*a*_|*data*) = *P*(⋃_*l*_ *β*_*jl*_ ≠ 0 |*data*), 1 ≤ *j* ≤ *p*. We then define the overall BFDR(55) as: 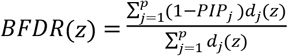, where *d*_*j*_(*z*) = *I*{*PIP*_*j*_ < *z*}. In the two real-data application examples, we use the overall BFDR to select putative causal genes.

FM-GPT is implemented as an R package with computationally efficient C++ codes integrated via Rcpp. The package takes individual-level genotype (within a region of interest) and phenotype data as input and output prioritized putative causal genes along with their PIP, BFDR and the phenotypes factors they target at. The package is freely available at www.github.com/tacanida/fm-gpt.

### 5. Comparative methods

We compared our proposed method to several existing fine-mapping approaches. For TWAS fine-mapping methods, we primarily compared against GIFT(14), a frequentist method that focuses on conditional gene effects and is designed for single-trait analysis. GIFT can be implemented using either a one-stage or two-stage framework, with the two-stage GIFT being equivalent to MVIWAS(21). We also attempted to implement FOCUS (15) and FOGS(16), but both methods were computationally infeasible and did not complete within a reasonable time. mvSuSIE (23) is an approximate Bayesian method that decomposes SNP effects into “single effects” to identify causal SNPs, and it is the only fine-mapping method designed to simultaneously target multiple traits. PAINTOR (17) and CAVIAR (22) are frequentist methods primarily developed for GWAS fine-mapping that directly model SNP linkage disequilibrium (LD) to identify causal variants. For single-trait methods (e.g., GIFT), we applied factor analysis first and then implemented the method using either single or multiple factors in simulations, or applied them separately to each trait in real data example 2. Detailed implementation procedures and parameter settings are provided in the Supplement S2.

For Bayesian fine-mapping methods, including our own, we controlled for false discoveries by applying Bayesian false discovery rate (BFDR) procedures to select genes. For frequentist fine-mapping methods that produce p-values, we applied the Benjamini-Hochberg (BH) procedure(59) to control the FDR when selecting genes.

### 6. Description of database and cohort used in the real data application

#### The Genotype-Tissue Expression (GTEx) project

The Genotype-Tissue Expression (GTEx) project(1) is a comprehensive resource designed to study the relationship between genetic variation and gene expression, by collecting postmortem tissue samples from over n_1_=800 deceased donors, covering approximately 50 distinct human tissue types. This extensive collection enables the characterization of tissue-specific regulatory effects of genetic variants, particularly expression quantitative trait loci (eQTLs), which are critical for understanding the genetic architecture of complex traits and diseases and have been widely used for TWAS(3). For our analyses, we utilized the predicted gene expression weights generated by the GTEx Consortium (v8 release), available through the PredictDB repository (https://predictdb.org/post/2021/07/21/gtex-v8-models-on-eqtl-and-sqtl/). These pre-trained models cover multiple tissues and provide tissue-specific weights essential for TWAS.

#### UK Biobank (UKB)

The UK Biobank (UKB) is a large-scale prospective cohort study that recruited over 500,000 participants aged 37–73 years between 2006 and 2010 across 22 assessment centers throughout the UK ((60)). At baseline, participants provided extensive lifestyle, medical, and demographic information, as well as biological samples for genotyping, creating a rich resource for studying the genetic and environmental determinants of complex traits. For our study, UKB provides critical resources for two complementary applications. In the first example, structural brain MRI data collected for approximately ∼30k participants starting in 2014 allows us to quantify cortical thickness measures across the whole brain. These high-resolution imaging phenotypes, combined with genetic data, enable the investigation of the genetic architecture of brain structure. In the second example, UKB’s extensive EHR data capture a wide range of disease diagnoses, treatments, and laboratory measurements across multiple organ systems. By linking these EHR-derived phenotypes with genotypes, we can perform phenome-wide analyses to identify causal genes influencing multiple related but heterogeneous disease conditions.

## Acknowledgments

Research reported in this publication was supported by the National Institute on Drug Abuse (NIDA) of the National Institutes of Health under award number 1K01DA059603-01A1, by the University of Maryland Grand Challenge Grant and EPIB Pilot Award to TM. This research was conducted using the UK Biobank Resource under application 74376. The content is solely the responsibility of the authors and does not necessarily represent the official views of the National Institutes of Health.

## Contributors

TC and ZY contributed equally as co-first authors. Concept and design: TC and TM. Method development: TC, SW, HH and TM. Simulation conduction: TC and ZY. Acquisition, analysis, or interpretation of real data: TC, ZY, YP, ML and TM. Drafting of the manuscript: TC, ZY and TM. Software development: TC. Obtained funding: SC and TM. Supervision: TM. All authors provided critical feedback on the research, analysis, and manuscript, and approved the final version.

## Notes

### Competing Interest Statement

The authors have declared no competing interest.

